# PhenoExam: an R package and Web application for the examination of phenotypes linked to genes and gene sets

**DOI:** 10.1101/2021.06.29.450324

**Authors:** Alejandro Cisterna, Aurora González-Vidal, Daniel Ruiz, Jordi Ortiz, Alicia Gómez-Pascual, Zhongbo Chen, Mike Nalls, Faraz Faghri, John Hardy, Irene Díez, Paolo Maietta, Sara Álvarez, Mina Ryten, Juan A. Botía

## Abstract

Gene set based phenotype enrichment analysis (detecting phenotypic terms that emerge as significant in a set of genes) can improve the rate of genetic diagnoses amongst other research purposes. To facilitate diverse phenotype analysis, we developed PhenoExam, a freely available R package for tool developers and a web interface for users, which performs: (1) phenotype and disease enrichment analysis on a gene set; (2) measures statistically significant phenotype similarities between gene sets and (3) detects significant differential phenotypes or disease terms across different databases. PhenoExam achieves these tasks by integrating databases or resources such as the HPO, MGD, CRISPRbrain, CTD, ClinGen, CGI, OrphaNET, UniProt, PsyGeNET, and Genomics England Panel App. PhenoExam accepts both human and mouse genes as input. We developed PhenoExam to assist a variety of users, including clinicians, computational biologists and geneticists. It can be used to support the validation of new gene-to-disease discoveries, and in the detection of differential phenotypes between two gene sets (a phenotype linked to one of the gene set but no to the other) that are useful for differential diagnosis and to improve genetic panels. We validated PhenoExam performance through simulations and its application to real cases. We demonstrate that PhenoExam is effective in distinguishing gene sets or Mendelian diseases with very similar phenotypes through projecting the disease-causing genes into their annotation-based phenotypic spaces. We also tested the tool with early onset Parkinson’s disease and dystonia genes, to show phenotype-level similarities but also potentially interesting differences. More specifically, we used PhenoExam to validate computationally predicted new genes potentially associated with epilepsy. Therefore, PhenoExam effectively discovers links between phenotypic terms across annotation databases through effective integration. The R package is available at https://github.com/alexcis95/PhenoExam and the Web tool is accessible at https://snca.atica.um.es/PhenoExamWeb/.

## Introduction

One of the main aims of clinical genetics research is to discover new gene-disease associations [1–6]. A disease is commonly diagnosed through the identification of a set of symptoms and signs associated with a particular and recognized clinical phenotype [8–10]. While some phenotypes are due to the impact of environmental factors, if a disease is inherited then the genetic variation within the individual also explains the phenotype at least partially [11]. Here, we introduce PhenoExam, a software tool to assist in the identification of new gene-phenotype associations. PhenoExam focuses on genetic diseases, harnessing all available gene-phenotype annotation resources to provide a comprehensive gene set and differential gene set annotation approach.

Over the last decade, we have seen attempts to standardize our knowledge of genetic diseases by formally linking genes to phenotypes using standard terminology, as exemplified by The Human Phenotype Ontology (HPO) [12] and The Mouse Genome Database (MGD) [13]. HPO is a standardized set of human phenotypic terms that are organized hierarchically with a directed acyclic graph and have been used to annotate all clinical entries in the Online Mendelian Inheritance in Man database (OMIM). OMIM [14] is a continuously updated catalog of human genes, genetic diseases and traits, with a particular focus on the molecular relationship between genetic and phenotypic variation. On the other hand, MGD is the manually curated consensus representation of genotype to phenotype information including detailed information about genes and gene products. It is the authoritative source for biological reference data sets related to mouse genes, gene functions, phenotypes, and mouse models of human disease. MGD has more terms and detailed phenotypic information than HPO because scientists can perform a wider set of experiments on mice. These features increase our knowledge and can help to prioritize novel gene-phenotype relationships in humans. Beyond phenotype databases, PhenoExam also includes gene-disease association databases, namely UniProt [15], The Comparative Toxicogenomics Database (CTD) [16], Orphanet [17], The Clinical Genome Resource (ClinGen) [18], The Genomics England PanelApp [19], The Cancer Genome Interpreter (CGI) [20] and PsyGeNET [21]. It also includes CRISPRbrain [22], the first genome-wide CRISPR interference and CRISPR activation screen in human neurons so we may study the potential association of phenotypic terms to specific functions of these genes in human neurons.

Apart from being a general-purpose tool for phenotype-based gene sets annotation, PhenoExam can also help in the diagnosis of genetic diseases. Currently fewer than half of patients with suspected Mendelian disorders (genetic diseases primarily resulting due to alterations in one gene) receive a molecular diagnosis [23]. Diseases with a genetic basis are usually diagnosed by looking for causal mutations in a panel of genes specifically associated with the disease. Gathering all phenotypes associated with the genes in a panel delivers a general phenotype-level description beyond the disease under study. To improve the accuracy of genetic diagnosis, we need methods to appropriately evaluate the gene level phenotypic similarity between candidate diseases. Moreover, the identification of differential phenotypes between diseases can also help towards more precise diagnostics. The identification of exclusive and/or shared phenotypes between gene panels can demonstrate common pathophysiology [24] but it can also help to create genetic links between diseases through their gene sets [25–26]. We can find numerous methods based on measuring disease-based phenotypic similarities by comparing sets of HPO terms e.g., Phenomizer [27], HPOSim [28], and PhenoSimWeb [29], Table 1 offers a detailed comparison amongst all tools. We also have modPhEA [30], an online resource for phenotype enrichment analysis. modPheEA helps with the gene-based phenotype enrichment analysis but just focused on one phenotype database at a time and without considering conditional analyses (two gene sets).

**Table 1.** Comparison of PhenoExam and other similar tools. “X” means the tool provides the function and “−” means the tool does not. “*” means the similarity scores are between phenotype terms and not between gene sets as does PhenoExam.

| Tool | As web | As software tool | Open source | Model Organism | Phenotype sets | Gene sets | Multiple database at once | Phenotype Enrichment Analysis | Disease Enrichment Analysis | Differential phenotypes | Diagnosis based on phenotypes | Similarity scores |
| --- | --- | --- | --- | --- | --- | --- | --- | --- | --- | --- | --- | --- |
| PhenoExam | X | X | X | X | X | X | X | X | X | X | - | X |
| modPhEA | X | - | - | X | X | X | - | X | - | X | - | - |
| DisGeNET | X | X | X | - | X | X | X | - | X | - | - | - |
| Phenomizer | X | - | - | - | X | - | - | - | - | - | X | * |
| HPOSim | - | X | X | - | X | X | - | X | - | - | - | * |
| PhenoSimWeb | X | - | - | - | X | X | - | - | - | - | - | * |

Phenomizer obtains the phenotype semantic similarity between sets of phenotypes based on the HPO ontology but does not rely on the use of the genes implicated in each phenotype. HPOSim is an R package that implements widely used ontology-based semantic similarity measurements to quantify phenotype similarities, and phenotype-level enrichment analysis using a hypergeometric test and the NOA method [31]. PhenoSimWeb is an online tool for measuring and visualizing phenotype similarities using HPO, uses a path-constrained Information Content-based measurement in three steps and exploits the PageRank algorithm [32]. Nevertheless, these tools did not take some important concepts into consideration.

PhenoExam contributes to the field with new features. These include the ability to detect differential phenotypes between pairs of gene sets: phenotypes that are significant within one gene set only, useful for detecting featured phenotypic terms between gene sets to distinguish better between similar diseases. It also combines phenotype and disease terms. This is important to link phenotypes to specific diseases. Finally, it tries to make the interpretation of the results of the phenotypic analysis easier by using simple scores to rank significant terms as well as summary messages and interactive graphs. We also found a knowledge management platform integrating and standardizing data about disease-associated genes from multiple sources called DisGeNET [33]. While being similar to PhenoExam in finding gene-disease associations, DisGeNET does not, however, offer facilities for gene-based phenotype enrichment analysis or for detecting phenotypic similarities between pairs of gene sets. PhenoExam uses as the basic substrate for gene-phenotype and gene-disease associations a number of configurable databases both in human and mouse that the user can tailor and adapt depending on the type of analysis to be performed. In PhenoExam, the phenotypic similarity between two groups of genes is performed by assessing the statistical significance of the Phenotypic Overlap Ratio (POR) between those (i.e. the number of common enriched phenotypes between the gene sets) (See methods Phenotype scores calculation).

We developed PhenoExam intending to support a variety of target users, mainly clinicians, computational biologists and geneticists. PhenoExam can help clinicians with finding phenotypes which are exclusive to diseases amongst a set of possible genetic disease candidates whose diagnosis is based on gene sequencing panels (Case 1). PhenoExam is also useful for geneticists as it can be used to improve their in-house-maintained gene panels but also to more accurately select genes involved in specific genetic studies (Case 2). Finally, computational biologists can use PhenoExam to discover new information about gene sets of interest thanks to the integration of multiple phenotype and disease databases and to compare phenotypes between known genes associated with a disease and the validation of computationally predicted disease genes (Case 2).

## Design and implementation

### Database Integration

The set of analyses performed by PhenoExam is based on manually curated phenotypes language like HPO, gene-disease ones as OMIM but also screening-based databases like CRISPRBrain, amongst many others (see table 2 for a complete list, description, and potential use). PhenoExam can perform a variety of analyses (Figure 1). The integration of these different databases is possible thanks to a well-established standardization process of genes and phenotypes used by PhenoExam. This includes using the HGCN Gene Nomenclature Committee (HGCN) gene naming system as the common way of identifying all human genes, and the definition of a new annotation term within each annotation database to indicate the HGCN genes that do not have any phenotype term associated in the database of interest. The list of HGCN genes was obtained from [34] https://www.genenames.org/download/statistics-and-files/. The HPO gene-phenotype association list was obtained from https://archive.monarchinitiative.org/latest/tsv/gene_associations/. The new no-phenotype association (HPO:XXX No HPO phenotype) was added to HPO for all protein coding genes with no known association to phenotype. For MGD, MP terms from orthologous genes to humans were obtained from http://www.informatics.jax.org/downloads/reports/index.html#go, and the relationship between human genes - mouse genes - mouse phenotype were collected using the files (MGI_PhenoGenoMP.rpt, HMD_HumanPhenotype.rpt, VOC_MammalianPhenotype.rpt). A new no- phenotype association (MP:XXX No phenotype) was created and all the protein coding genes without a relation to phenotype were linked to this term. For CRISPRBrain, the gene-phenotype relationships were obtained from https://crisprbrain.org/simple-screen/. For the generation of this database, the phenotypes were codified in three classes for each CRISPR analysis: association to the phenotype (Positive-Hit and Negative-Hit genes in CRISPRBrain), positive association (Positive-Hit genes in CRISPRBrain) and negative association (Negative-Hit genes in CRISPRBrain). This was accomplished according to the Hit-Class label in CRISPRbrain (Positive-Hit, Negative-Hit). The non-relationship phenotype (CRB:XXX No phenotype was created and all the protein coding genes that were not related to any phenotype were related to this term. We integrate into PhenoExam only the information from curated databases (UniProt, CTD, Orphanet, ClinGen, The Genomics England PanelApp, CGI and PsyGeNET). Then the non-relationship disease term (CXXX No diseases associated) was created and all the protein coding genes that were not related to any disease were related to this term. After standardization process, the current release (v1.0) of PhenoExam contains, 659634 gene-phenotype associations, involving 20209 genes, 18159 different phenotypes and 9348 different diseases (see details in Table 2).

**Table 2.** Databases usable through PhenoExam and size of each in terms of genes, phenotypes and associations. Numbers reported are final, after preprocessing and unification of gene names across databases.

| Source | Genes | Phenotypes | Diseases | Assocs | Summary |
| --- | --- | --- | --- | --- | --- |
| HGCN | 19197 | - | - | - | All protein coding genes |
| HPO | 19248 | 7861 | - | 186290 | Human gene-phenotype associations |
| MGD | 17900 | 10243 | - | 242313 | Mouse gene-phenotype associations |
| CRISPRBrain | 19275 | 55 | - | 43481 | Cell screen gene-phenotype associations |
| ClinGen | 19198 | - | 420 | 19851 | Human gene-disease associations |
| Genomics England | 19230 | - | 5538 | 24336 | Human gene-disease associations |
| CTD | 19636 | - | 6843 | 58660 | Human gene-disease associations |
| CGI | 19198 | - | 177 | 20361 | Human gene-disease (cancer) associations |
| UniProt | 19204 | - | 3868 | 21101 | Human gene-disease associations |
| Orphanet | 19262 | - | 3183 | 2228 | Human gene-disease (rare) associations |
| PsyGeNET | 19248 | - | 82 | 20952 | Human gene-disease associations |
| **ALL** | **20209** | **18159** | **9348** | **544022** | **PhenoExam tool** |

**Figure 1.**
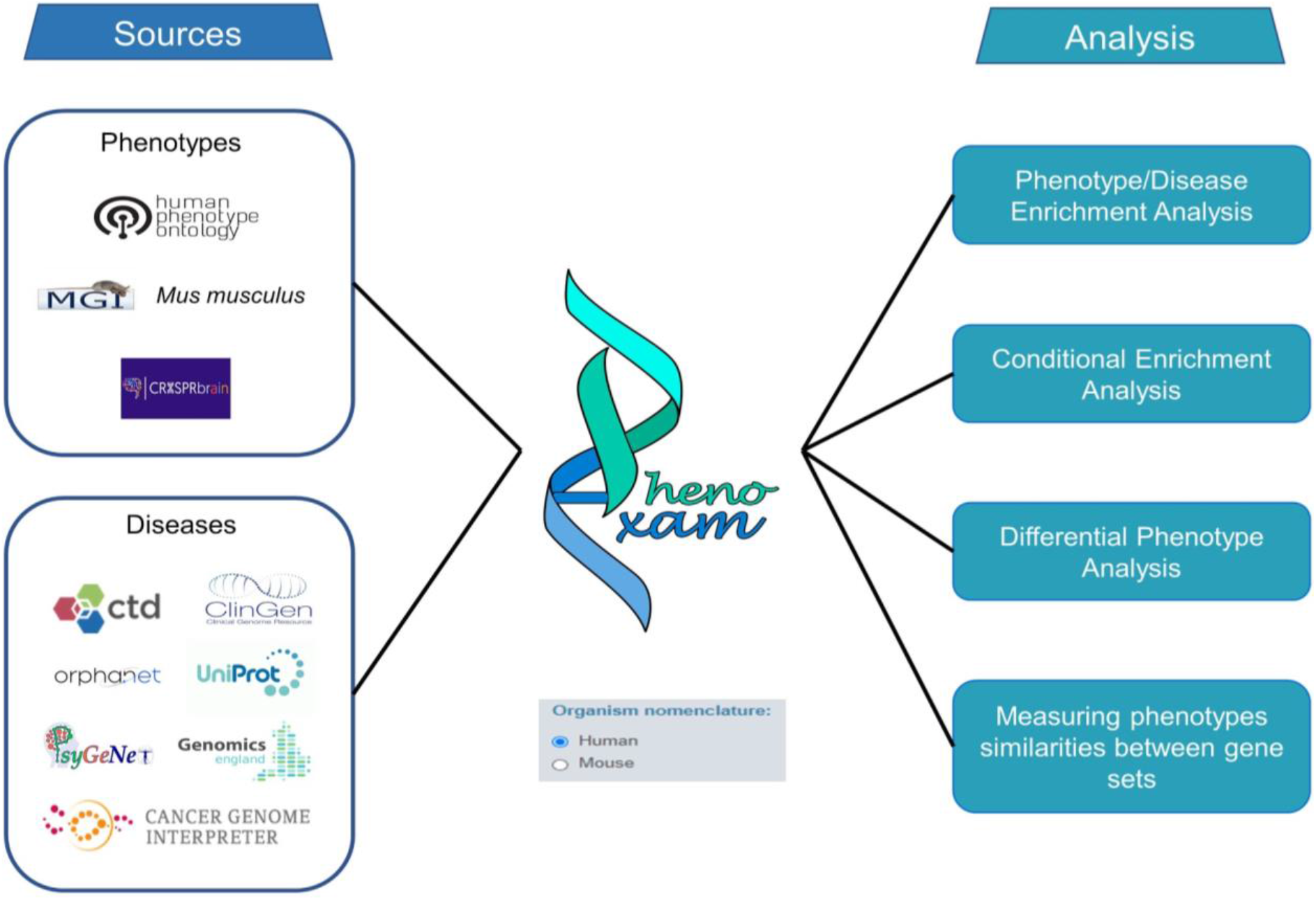
Schematic representation of PhenoExam integrated databases and offered analyses. We can use PhenoExam with human or mouse genes. PhenoExam annotation databases include HPO, MGI, CRISPRBrain, CTD, ClinGen, OrphaNET, UniProtm PsyGeNET, CGI and Genomics England. The tool offers a variety of analyses. Given a gene set of interest, G, the user can evaluate its enrichment for phenotypes and disease in all or a subset of the offered databases. Given two gene sets, G and G’, the user can evaluate whether the phenotype terms enriched in G are also enriched in G’ when G and G’ do not overlap e.g., G’ was predicted from G, with the Conditional Enrichment Analysis. If G and G’ show some gene overlap, the user can assess whether the gene sets show any differential phenotypes through the Differential Phenotype Analysis.

### Phenotype scores calculation

#### Phenotype Enrichment Analysis on a gene set G

PhenoExam obtains a list of statistically significant enriched phenotypes in a given set of gene *G* within a phenotype/disease database annotation of reference D. In order to calculate whether a gene set *G* shows enrichment in a given phenotypic term *p* belonging to D, let g be the number of genes in G associated with p. Let also gdb be the number of genes associated with p and GDB the total number of genes in the database, we model the enrichment probability with a hypergeometric distribution such that:

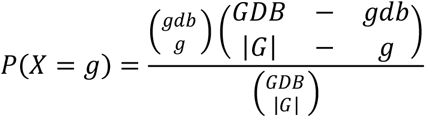

Any phenotype with *p < 0.05* will be enriched in the *G* gene set. We compute this probability for each phenotypic term *ph* associated with 1 gene or more in *G*, and use these probabilities as p-values. PhenoExamWeb reports the raw and Bonferroni adjusted p.values.

#### Phenotypic Overlap Ratio score

PhenoExam’s approach to measuring the similarity between two gene sets G and G’, within an annotation database D, is based on a score called the Phenotypic Overlap Ratio (POR). Let Gp be the number of significantly enriched terms in D for genes in G, and analogously for G’p. The POR is the Jaccard index on the agreement between the subsets of significant phenotypes:

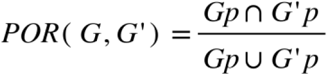

POR (*G,G’*) takes values in [0,1], resulting in 0 when no phenotype is shared and 1 when the sets share all phenotypes.

#### Statistically significant Phenotypic Overlap Ratio

PhenoExam assess whether the POR between gene sets G and G’ is statistically significant by means of randomization. We will have two modalities of the POR, depending on whether G and G’ share genes or, on the contrary, they are disjunct (e.g., G’ was predicted from G). When G and G’ are thought to share genes, POR (*G,G’*) is compared with POR (*G,R*) and with POR (*G’,R’*), where *R* has the same size as G and R’ the same as G’. Genes in both R and R’ are chosen randomly within the whole set of protein coding genes. We repeat this process for m random gene sets *R*_*1*_, *R*_*2*_,…, *R*_*m*_ and *R^′^_1_*, *R^′^_2_*,…, *R^′^_m_* to obtain an empirical p-value with the proportion of random gene sets whose POR is greater than the observed one. On the other hand, when G’ is obtained by using G as input of the generation process, we say G’ is conditioned to G. Therefore, the significance test of the POR(G, G’) is reduced now to obtain an empirical p-value based on the proportion of times a randomized POR(G,R), with R any of *R*_*1*_, *R*_*2*_,…, *R*_*m*_ all with the same size of G while keeping G constant, shows higher values than the observed POR(G, G’).

#### Relaxed Phenotypic Overlap Ratio

The POR only considers phenotypes that were assessed as statistically significant. Sometimes, it may be of interest to relax this restriction to incorporate all phenotype/disease terms associated with G. In this case, the score is called Relaxed Phenotypic Overlap Ratio (RPOR). It is calculated in a similar way to the POR but with all phenotypes, whether these are enriched or not. In the same way, as with the POR, we can determine whether the RPOR is statistically significant by using randomization.

#### Phenotype relevance association analysis for gene sets

Once it has been determined that two sets of G and G’ genes share some enrichment of phenotypic terms, and focusing only on the shared terms, we can measure the correlation of the number of genes of each phenotypic term as measured in G and G’ by a linear regression model and report the R^2^ as the strength of this correlation together with the association p-value. Higher values of R^2^ would suggest a linear association between importance of phenotypic terms in G and importance of the same genes in G’.

### Generation of the web interface

We have developed PhenoExamWeb, a web based tool for performing phenotypic analyses using R. PhenoExamWeb shiny app is accessible at https://snca.atica.um.es/PhenoExamWeb/. R and shiny R package [35] were used for front-end scripting of the web interface. R script was used for back-end execution and analysis with the development environment of R version 3.6.3. The R package is available at https://github.com/alexcis95/PhenoExam.

### Analysis with PhenoExamWeb

PhenoExamWeb requires gen symbols, human or mouse, as the input file. Then, we need to select the type of analysis: Phenotype Enrichment Analysis (One gene set) or Phenotype Comparator (Two gene sets). We also need to specify the database or databases. The workflow of PhenoExamWeb is summarized in Figure 2. Users can follow the web tutorial on the website (https://snca.atica.um.es/PhenoExamWeb/#section-help) and the R package tutorial on GitHub (https://rawcdn.githack.com/alexcis95/PhenoExamWebTutorials/b3d40397e0f41af5ff50462350fdc81369479810/tutorial.html).

**Figure 2.**
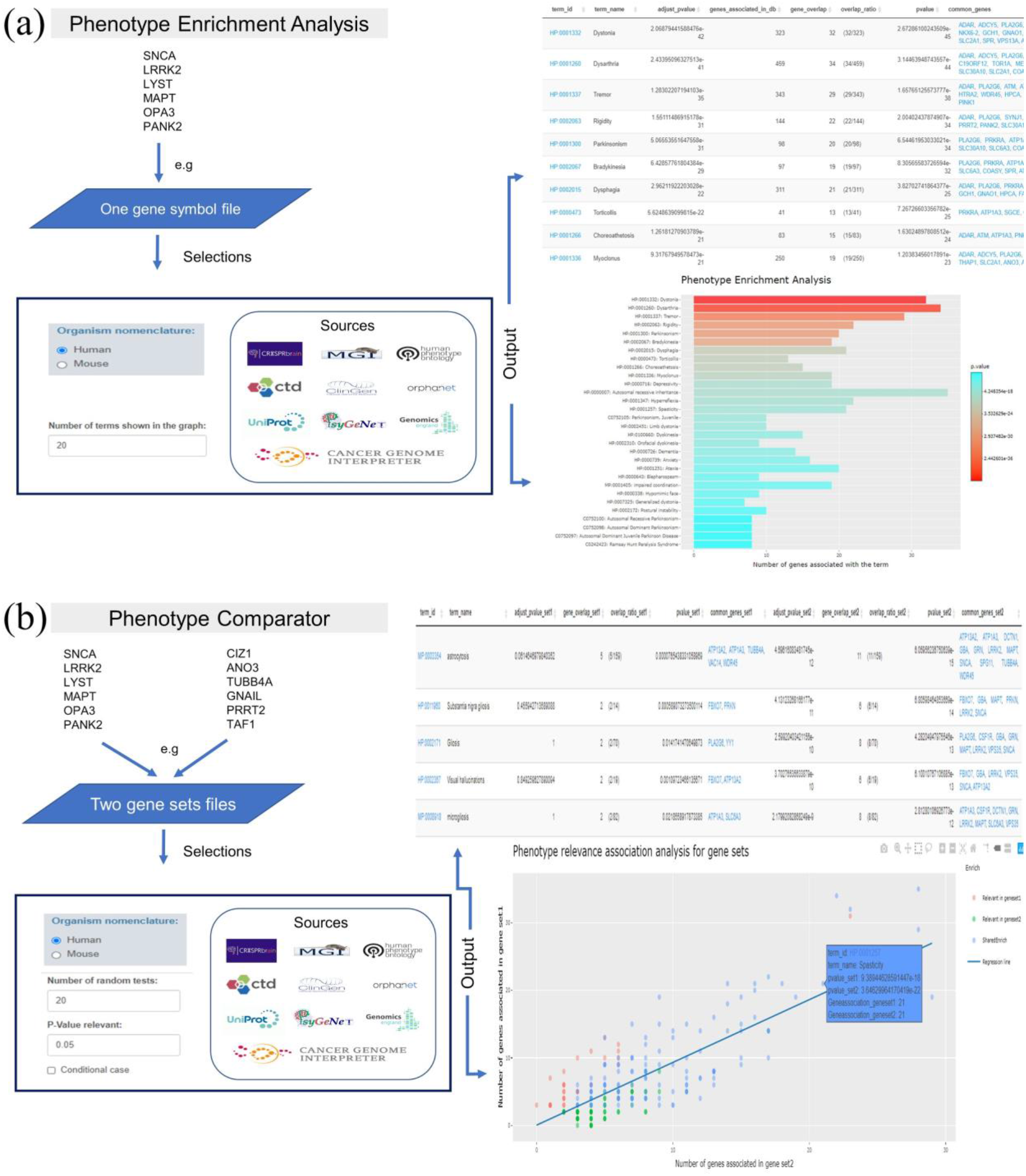
PhenoExamWeb shiny app possible workflows. (a) Phenotype Enrichment Analysis: requires one gene symbol file as input file, which gene symbol nomenclature (Organism nomenclature: Human or Mouse) we use, the phenotype/disease annotation databases to be considered and the top number of terms shown in the graph. The results generate an interactive table and graph which include phenotypes, genes implicated with each term and p.values as output. (b) The Phenotype Comparator requires two gene sets as input together with the gene symbol nomenclature (Human or Mouse) used, the annotation databases of interest for the analysis and the number of random tests to obtain empirical p-values, the relevant p-value threshold and whether our analysis is a conditional case (i.e. if one gene set was generated after a prediction analysis from the other and they are totally different gene sets). Finally, we obtain the summary of the analysis with the similarities phenotype scores, the differential phenotypes, interactive tables and graphs with phenotypes, genes and p.values as output for detailed inspection and result presentation.

## Results and discussions

### PhenoExam controls Type I error when used with all phenotype databases

We assessed PhenoExam for type I error given all phenotype/disease databases considered in the task of phenotypic enrichment analysis of gene sets. Firstly, we evaluated the possibility of finding a phenotypic term erroneously enriched, due to random chance, amongst all the terms at the database, for gene sets of varying sizes. For such purpose we performed simulations of phenotype enrichment analysis for different random gene sets with a variable number of protein coding gene sizes (5, 10, 20, 40, 80, 160, 320, 640) tested in all annotation databases. Each combination of gene set size and database was simulated 1000 times, yielding a total of 80000 simulations. A graphical representation of the summary of results appears in Figure 3. PhenoExam maintains Type I error under control, see figure 3, (a) plot, with a significance level of 0.05 as the number of significant tests is always under 0.05 ratio. We observed a negative correlation between gene set size and proportion of false positive tests, r = −0.453, p = 0.026. We found a different tendency on the Type I error with some disease databases, specially Genomics England Panel App (GEL) and Orphanet. With those, PhenoExam only controls Type I error when the gene set size is high (greater than 80 for Orphanet, 160 for Genomics England). The rest of databases are usable with a gene set size over 40. The main reason is the low average number of disease terms associated with each gene, 4.39 for GEL and 7 for Orphanet, in comparison with the average for the rest of disease databases, 17.7. We also found, as we expected, a negative correlation between the number of genes per random set and the type I error, r = −0.381, p = 0.0038. For these reasons, we recommend prioritizing the selection of the CTD database for disease analyses when we have less than 40 genes. Users can find more information about what database they need to use at https://snca.atica.um.es/PhenoExamWeb/#section-help

**Figure 3.**
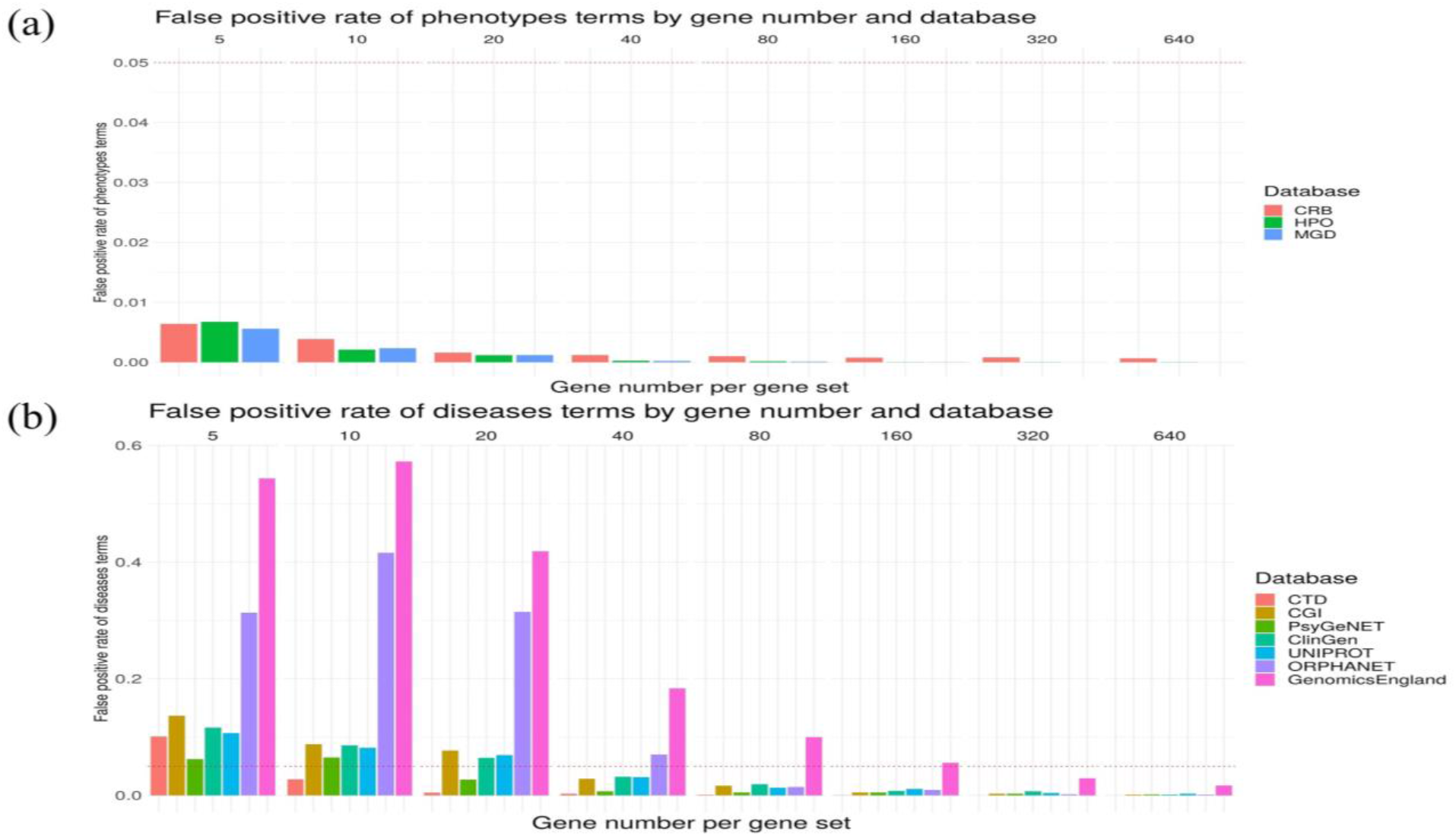
False positive rate of phenotype (a) and disease (b) terms enrichment across varying gene set sizes (5, 10, 20, 40, 80, 160, 320, 640) per phenotype/disease database. As the simulation points out, CRB, HPO, MGD, are perfectly usable for any gene set size, CTD is recommended for gen set sizes over 10, PsyGeNET for 20, CGI, ClinGen and Uniprot for 40, Orphanet for 80 and GEL for gene set sizes over 160.

### PhenoExam differentiates between gene sets with very similar phenotypes

We evaluated how accurate PhenoExam is when computing the POR (detecting phenotype similarities) between gene sets by comparing genetic forms of epilepsy (261 genes from NIMGenetics epilepsy panel) and “artificial” gene sets constructed with variable POR with the original epilepsy gene set and additional genes with similar phenotypic connectivity not associated to epilepsy. In these additional genes we injected a 5% of noise with genes associated with epilepsy phenotypic terms. We performed 1000 simulations for the artificial genes sets (261 genes) constructed with different proportions of epilepsy genes between (0-100%) and different proportions of other genes (0-100%). We calculated the POR significance test between the real and the artificial gene sets (Figure 4). PhenoExam is sensitive in detecting differences between gene composition changes (≅1%) in different gene sets, which in this case are 3 genes. We observed a positive linear relationship between POR and the proportions of epilepsy genes in the artificial gene sets, 0.9674 R^2^ (P < < 2.2×10^−16^) (Figure 4a). We assessed that PhenoExam can distinguish well amongst the epilepsy real genes and the artificial gene sets constructed with high proportions of epilepsy genes (94-99% epilepsy genes) that gather very similar phenotypes with a t-test in all cases (P < 2.2×10^−16^) (Figure 4b).

**Figure 4.**
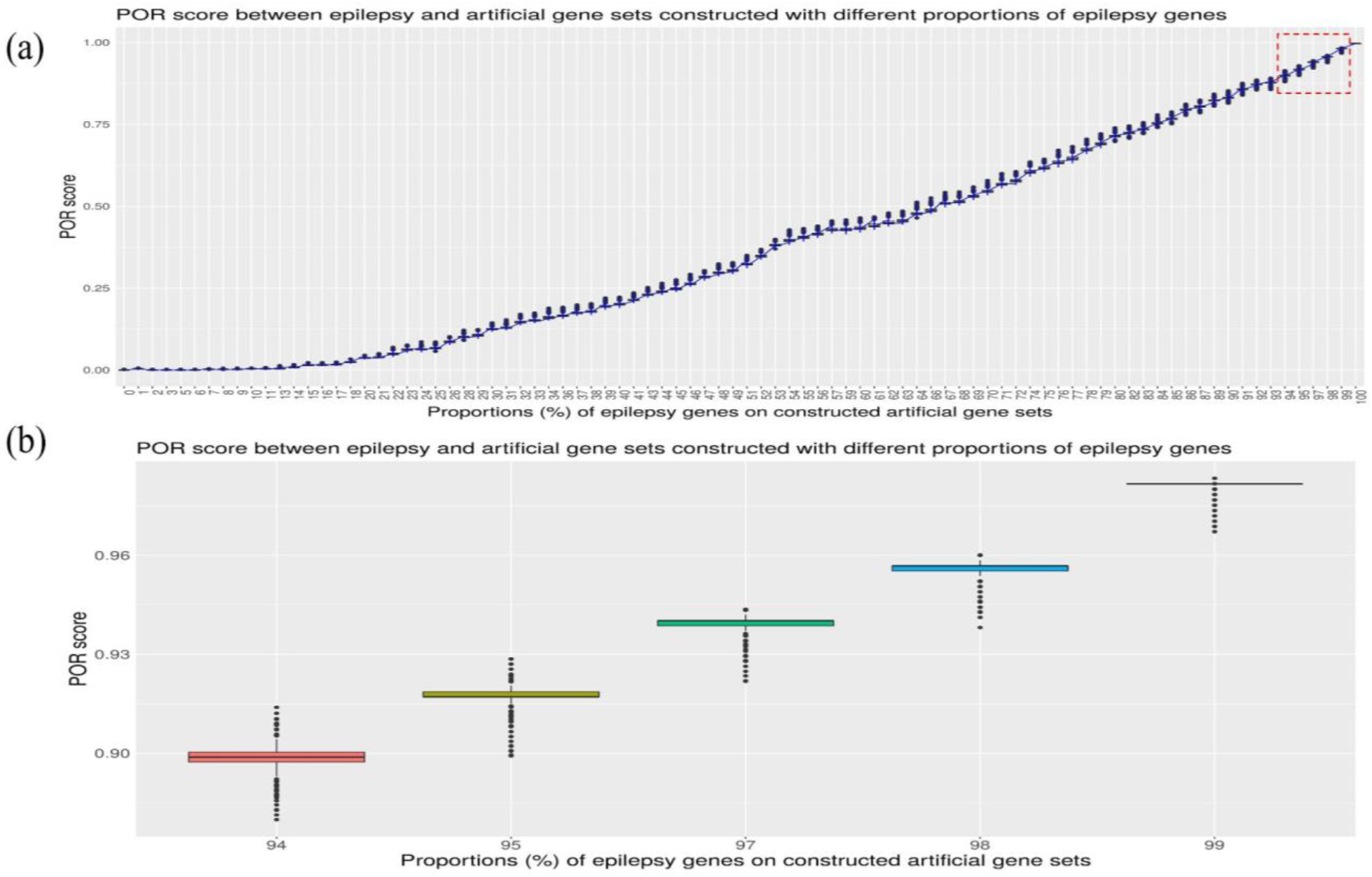
POR significance test between the real and the artificial gene sets constructed with different proportions of epilepsy genes (a) and detailed zoom of POR score between the real and the artificial gene sets constructed with different proportions of epilepsy genes (94-99% epilepsy genes). (a) We observed a positive linear relationship between POR and the proportions of epilepsy genes in the artificial gene sets, 0.9674 R^2^ (P < 2.2×10^−16^). (b) PhenoExam can distinguish well amongst the epilepsy real gene set and the artificial gene sets constructed with high proportions of epilepsy genes (94-99% epilepsy genes) that gather very similar phenotypes with a t-test in all cases (P < 2.2×10^−16^).

### Case 1: The analysis between juvenile-onset Parkinson’s disease (PD) and early onset dystonia (EOD) reveals they hold phenotype-level similarities but also potentially interesting differential phenotypes

We applied PhenoExam to the detection of differential phenotypes between gene sets by comparing two genetic diseases with similar symptoms: juvenile-onset Parkinson’s disease (PD) and early-onset dystonia (EOD). PD and EOD both are movement disorders, PD is caused by a degeneration in the basal ganglia and it has predominant symptoms consisting of tremor, rigidity, bradykinesia, postural instability and progressive dementia [36]. EOD is a disease characterized by involuntary muscle contractions leading to abnormal posturing and movements and postures, occurring with or without other neurological symptoms [37]. In our case we compared 35 PD genes and 50 EOD genes from Genomics England PanelApp (Supplementary file with genes “G1”), with 19 genes in the overlapping set (54.3% of genes on PD gene set). We ran a separate phenotype enrichment analysis for PD and EOD, using HPO, MGD, CTD and CRISPRBrain databases simultaneously (given the simulation analyses performed above, these are the databases recommended by PhenoExam) (Figure 5). We obtained an interactive table and graph with the enrichment phenotypes for PD and EOD (Supplementary Tables T1PD and T1EOD and Figures FS1PD and FS1EOD). The top two most enriched phenotypes, in each input database, for PD genes were Bradykinesia (HP:0002067; P = 2.16×10^−60^) and Parkinsonism (HP:0001300; P = 2.62×10^−51^) for HPO, Abnormal gait (MP:0001406; P = 3.78×10^−13^) and Neuron degeneration (MP:0003224; P = 9.98×10^−13^) for MGD, Parkinsonism, Juvenile (C0752105; P = 7.49×10^−28^) and Ramsay Hunt Paralysis Syndrome (C0242423; P = 7.49×10^−28^) for CTD, and no enrichment found for CRISPRBrain. All the enrichment terms found are supported by the literature [38–41]. At the EOD analysis, we found Dystonia (HP:0001332; P = 3.51×10^−42^) and Dysarthria (HP:0001260; P = 5.38×10^−41^) for HPO, impaired coordination (MP:0001405; P = 7.4×10^−14^) and Abnormal gait (MP:0001406; P = 3.17×10^−10^) for MGD, Parkinsonism, Juvenile (C0752105; P = 7.4×10^−13^) and Ramsay Hunt Paralysis Syndrome (C0242423; P = 7.4×10^−13^) for CTD, and again no enriched term for CRISPRBrain. Above mentioned phenotype terms are associated with dystonia according to several articles [42–46].

**Figure 5.**
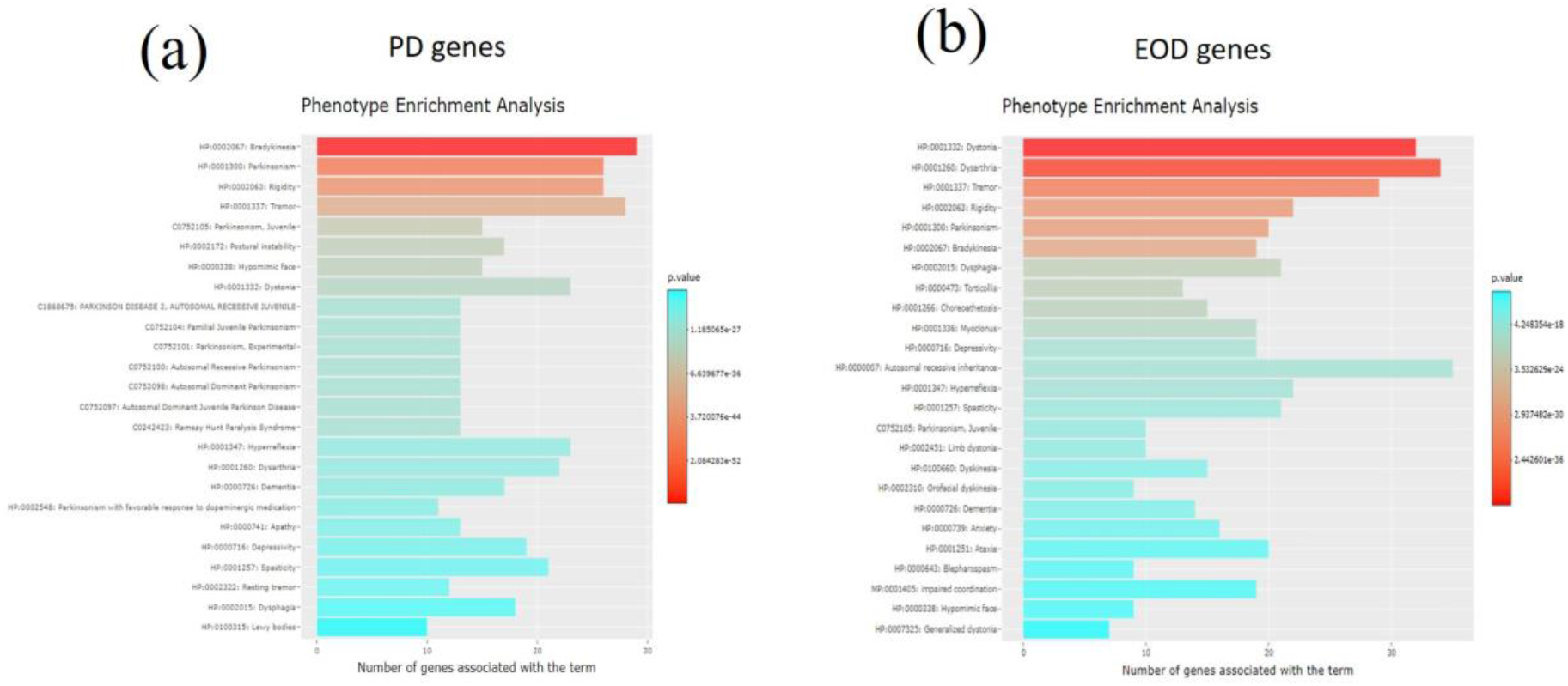
Phenotype Enrichment Analysis in PhenoExam for each gene set. The graph shows the 25 most enriched terms for PD genes (a) and for EOD genes (b).

We wanted to compare PD and EOD gene sets, through the Phenotype Comparator analysis in PhenoExamWeb (see figure 6) using HPO, MGD, CTD and CRISPRBrain as the databases selected, and a randomization based on 1000 null tests. This comparison yielded 139 shared significant phenotypic terms (out of 273 unique significant phenotypic terms in both, POR=0.509 (P < 0.001). Phenotype relevance association analysis for PD and EOD (i.e. whether the shared phenotypes are similar in relevance, i.e. in the number of genes associated with them, within each gene set) results in an adjusted R squared of 0.643 (P < 9.23×10^−63^) which suggests that an important portion of the common phenotypes are similar in relevance. We actually see they share phenotypic terms such as Tremor (HP:0001337), Bradykinesia (HP:0002067), Rigidity (HP:0002063), Dystonia (HP:0001332), Abnormal gait (MP:0001406) or Neuron degeneration (MP:0003224) (Supplementary Table T2Share). But we also detect differential phenotypes that can be displayed by interactive graphs and tables on the web. For example, significant terms exclusive from the PD gene set phenotypes include Astrocytosis (MP:0003354; P < 5.17×10^−12^), Substantia nigra gliosis (HP:0011960; P < 4.15×10^−11^), Neuronal loss in central nervous system (HP:0002529; P < 3.74×10^−6^), Orthostatic hypotension due to autonomic dysfunction (HP:0004926; P < 9.96×10^−6^) and Lewy Body Disease (C0752347; P < 1.11×10^−3^) (Supplementary Table T3PDdiff). Above mentioned phenotype terms are associated more or only with PD according to several articles [47–52]. The same analysis identified Writer’s cramp (HP:0002356; P < 1.37×10^−9^) as exclusive to EOD and this refers to a type of focal dystonia [53]. We also found Hypoplasia of the corpus callosum (HP:0002079; P < 3.56×10^−5^), a controversial and not widely studied phenotype in dystonia [54–55] and Acanthocytosis (HP:0001927; P < 2.76×10^−3^) a term normally associated with chorea-acanthocytosis, other disease with dystonia’s similar symptoms [56]. Microcephaly (HP:0000252; P < 4.17×10^−4^) is associated with dystonia and several genes such as KMT2B [57–58]. We also found Intellectual disability, mild (HP:0001256; P < 4.68×10^−3^), Dystonia, Primary (C0752203; P < 3.26×10^−7^) and Hyperactive deep tendon reflexes (HP:0006801; P < 4.31×10^−2^) that is associated with Paroxysmal dyskinesia (PxD) [59] (Supplementary Table T3EODdiff).

**Figure 6.**
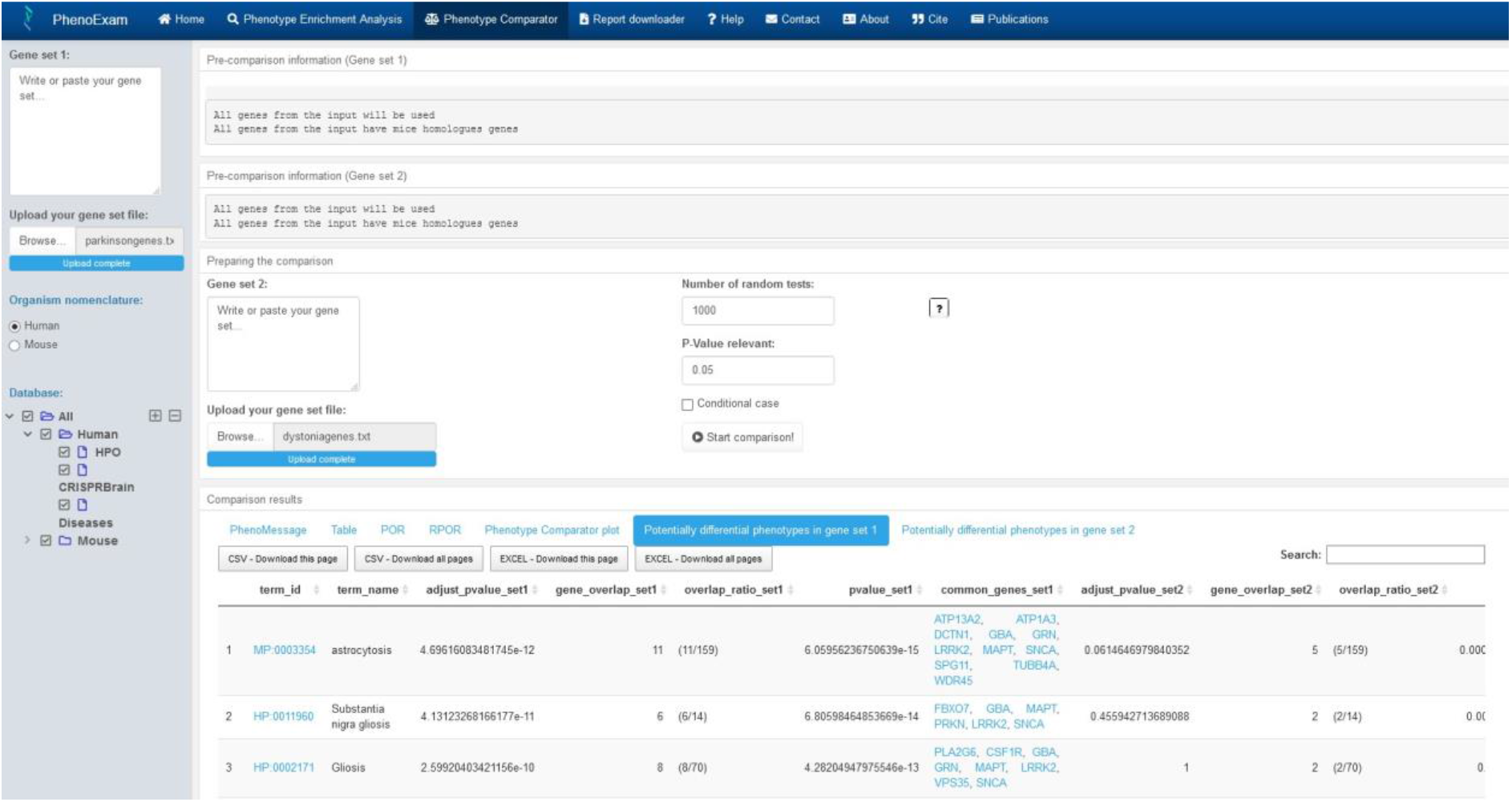
Phenotype Comparator analysis view. We selected PD genes as gene set 1, EOD genes as gene set 2, HPO, MGD, CRISPRBrain and CTD databases and 1000 random tests. We obtained as output interactive tables with the shared phenotypes and the differential phenotypes, plots, PhenoExam phenotype similarities scores and information.

### Case 2: New likely epilepsy genes predicted by G2PML recapitulate phenotype terms of known epilepsy genes

Let us suppose it is possible to discover new Mendelian genes associated with a specific disease (congenital epilepsy in this case) by finding non-linear patterns of the genes in that panel based on their description through properties based on genomic, transcriptomic and genetics of each gene with machine learning techniques. Therefore, in order to discover new genes, we aim at finding very similar genes in terms of those properties (see G2PML paper at biorxiv [60]). The question we face is: do those genes predicted to be linked to congenital genetic forms of epilepsy recapitulate similar phenotypes to the genes in the panel of origin? The more supportive the answer points to a phenotype recapitulation, the better the predictions made by G2PML. This is an example of what we call a conditional case, comparing phenotypes in gene sets G and G’ when they are disjunct and G’ was generated using G as seeds. More specifically, G refers to epilepsy genes from an in-house maintained epilepsy panel (261 genes) at NIMGenetics. Moreover, G’ is a set of 209 new genes as predicted by G2PML.

We carried out the Phenotype Comparator analysis in PhenoExamWeb with the conditional case option marked, gene set 1 was the epilepsy genes, gene set 2 was the new likely epilepsy genes predicted by G2PML, HPO, MGD, CRISPRBrain and CTD databases selected at the same time and we chose 1000 random tests. We obtained the Pheno Message from PhenoExamWeb that they shared 106 significant phenotypic terms (out of 734 unique significant phenotypic terms in both), which yields a POR of 0.144 (P < 0.001). Phenotype relevance association analysis for epilepsy associated genes and epilepsy predicted genes (i.e. whether the shared phenotypes are similar in relevance, i.e. in the number of genes associated with them, within each gene set) results in an adjusted R squared of 0.331 (P < 4.35×10^−66^) which suggests that an important portion of the common phenotypes are similar in relevance. The p-values were obtained through the randomization of 1000 random gene sets. We also obtained a table with the phenotypes shared between gene sets (Supplementary Table T4Share). New likely epilepsy genes predicted by G2PML, e.g. DDX3X, KCNH1, TBL1XR1, DLG4 or PDE2A, recapitulate phenotype terms of known epilepsy genes, we actually check they share epilepsy significant phenotypic terms such as Seizures (HP:0001250), Global developmental delay (HP:0001263), Microcephaly (HP:0000252), abnormal brain morphology (MP:0002152), hyperactivity (MP:0001399) and diseases terms without Bonferroni adjust Epilepsy (C0014544) and Autistic Disorder (C0004352). Above mentioned phenotype terms are associated with epilepsy according to several articles [61–69]. We also provided the number of genetic variants from the Epi25 whole-exome sequencing (WES) case-control study of each epilepsy gene predicted, we obtained 665 genetic variants in cases and 446 in controls (OR = 1.49)(Supplementary Table T5)[70].

## Conclusion

We developed PhenoExam, a freely available R package and Web application, which performs phenotype enrichment and disease enrichment analysis on gene set G, measures statistically significant phenotype similarities between pairs of gene sets G and G’ and detects statistically significant exclusive phenotypes or disease terms, across different databases. PhenoExam just required the names of genes in the gene sets as input and which databases to test for enrichment. It allows us to switch from the gene space and the phenotype space. PhenoExam can identify the statistically significant and differential phenotypes of a gene set as we showed with PD, EOD, epilepsy and likely epilepsy predicted genes. We proved with simulations that it is useful to distinguish between gene sets or diseases with very similar phenotypes through projecting genes into their annotation based phenotypical spaces. With the PD and EOD example above, we clearly see they hold phenotype-level similarities but also potentially interesting differential phenotypes. The conditional case studied between epilepsy associated and epilepsy predicted genes show they hold epilepsy phenotype terms in common, which is useful for the validation of computationally epilepsy predicted disease genes. Therefore, PhenoExam effectively discovers links between phenotypic terms across annotation databases by integrating different annotation databases. All these findings are supported with interactive plots (see tutorials at GitHub project) to foster the visualization and interpretation of findings.

## Supporting information

T1EOD

T2Share

T3EODdiff

T3PDdiff

T4Share

## Key points

- PhenoExam performs phenotypic analysis with gene symbols as input.
- PhenoExam is an R Package and is an easy to use freely available shiny app.
- PhenoExam integrates several databases such as HPO, MGD, CRISPRBrain, CTD, UniProt, Orphanet, ClinGen, Genomics England, CGI and PsyGeNET.
- Already tested with different sets of genes to prove the correct running of the tool.

## Availability and requirements

Project name: PhenoExam

Project home page: https://snca.atica.um.es/PhenoExamWeb/

Source code is available at https://github.com/alexcis95/PhenoExam

Operating system(s): Windows, Linux, Mac OS

Programming language: R language

License: GPL-2|GPL-3 Any restrictions to use by non-academics: none.

## Authors’ contributions

A.C., D.R. wrote the code; J.O. and A.G. tested the code. A.C. prepared the diagrams and figures. J.B., M.R. and A.C. wrote the manuscript with the help of the rest of authors. M.N., F.F., I.D., P.M, S.A. helped check and improve the manuscript. J.B. designed the study. All authors read and approved the final manuscript.

---

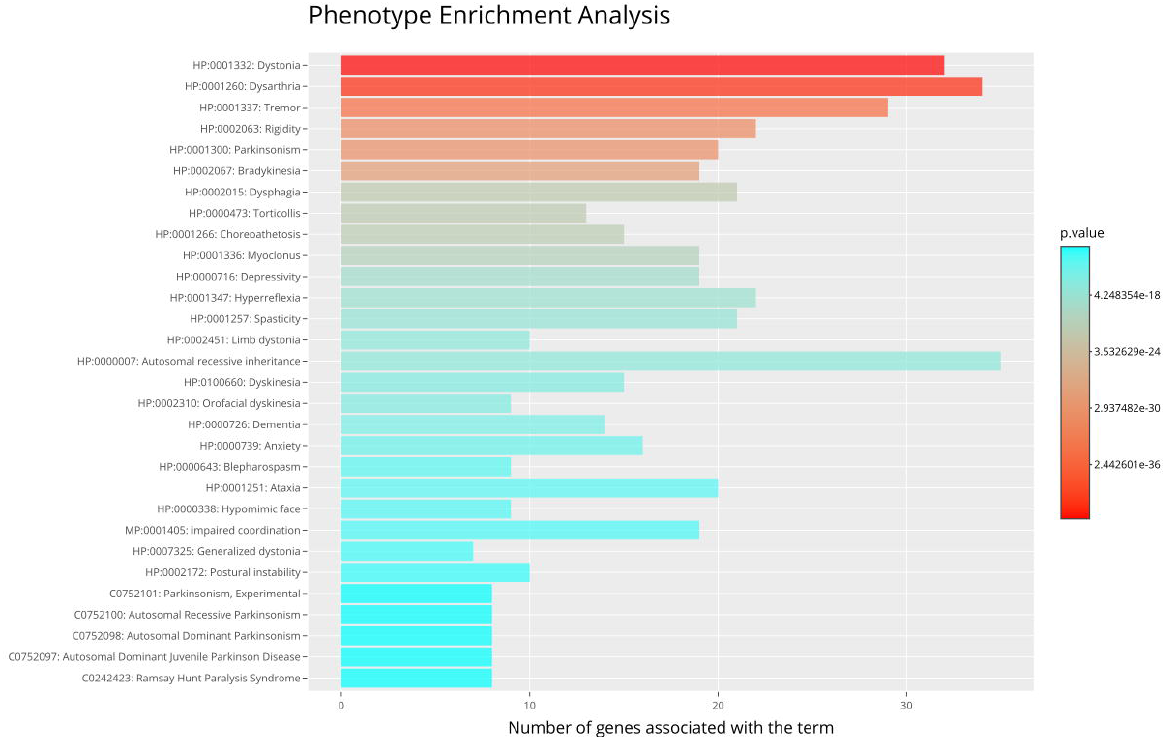

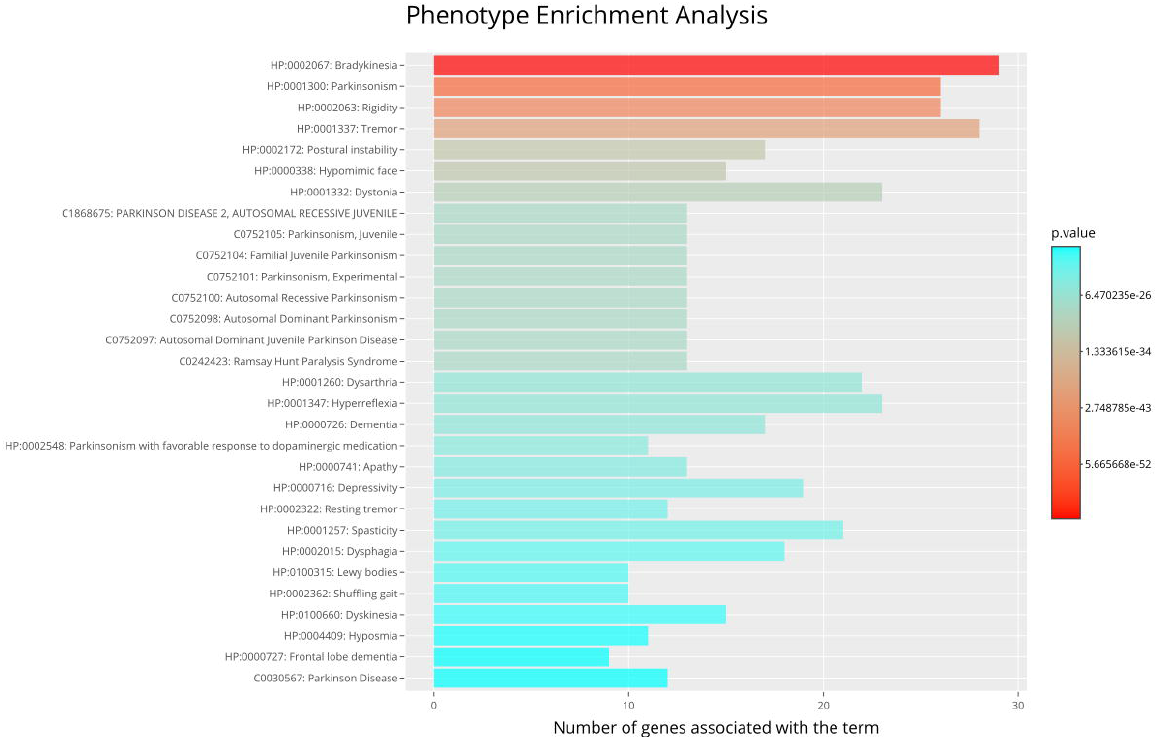

## References

[1] Jimenez-Sanchez, G., Childs, B., & Valle, D. (2001). Human disease genes. Nature, 409(6822), 853–855.

[2] Stankiewicz, P., & Lupski, J. R. (2010). Structural variation in the human genome and its role in disease. Annual review of medicine, 61, 437–455.

[3] Schaub, M. A., Boyle, A. P., Kundaje, A., Batzoglou, S., & Snyder, M. (2012). Linking disease associations with regulatory information in the human genome. Genome research, 22(9), 1748–1759.

[4] Shaw, C. J., & Lupski, J. R. (2004). Implications of human genome architecture for rearrangement-based disorders: the genomic basis of disease. Human molecular genetics, 13(suppl_1), R57–R64.

[5] Osborne, J. D., Flatow, J., Holko, M., Lin, S. M., Kibbe, W. A., Zhu, L. J., … & Chisholm, R. L. (2009). Annotating the human genome with Disease Ontology. BMC genomics, 10(S1), S6.

[6] Robinson, P. N., Köhler, S., Oellrich, A., Wang, K., Mungall, C. J., Lewis, S. E., … & Gilissen, C. (2014). Improved exome prioritization of disease genes through cross-species phenotype comparison. Genome research, 24(2), 340–348.

[7] Dorland, W. A. N. 1. (2012). Dorland’s illustrated medical dictionary. 32nd ed. Philadelphia: Elsevier/Saunders.

[8] Temple, L. K., McLeod, R. S., Gallinger, S., & Wright, J. G. (2001). Defining disease in the genomics era. Science, 293(5531), 807–808.

[9] Scully J. L. (2004). What is a disease?. EMBO reports, 5(7), 650–653. https://doi.org/10.1038/sj.embor.7400195

[10] Ritchie, M. D., Denny, J. C., Crawford, D. C., Ramirez, A. H., Weiner, J. B., Pulley, J. M., … & Haines, J. L. (2010). Robust replication of genotype-phenotype associations across multiple diseases in an electronic medical record. The American Journal of Human Genetics, 86(4), 560–572.

[11] Hunter, D. J. (2005). Gene–environment interactions in human diseases. Nature Reviews Genetics, 6(4), 287–298.

[12] Robinson PN, Köhler S, Bauer S, Seelow D, Horn D, Mundlos S. The human phenotype ontology: a tool for annotating and analyzing human hereditary disease. Am J Hum Genet. 2008; 83(5):610–5.

[13] Bult CJ, Blake JA, Smith CL, Kadin JA, Richardson JE, the Mouse Genome Database Group. 2019. Mouse Genome Database (MGD) 2019. Nucleic Acids Res. 2019 Jan. 8;47 (D1): D801–D806.

[14] Amberger, J. S., Bocchini, C. A., Schiettecatte, F., Scott, A. F., & Hamosh, A. (2015). OMIM.org: Online Mendelian Inheritance in Man (OMIM®), an online catalog of human genes and genetic disorders. Nucleic acids research, 43(Database issue), D789–D798. https://doi.org/10.1093/nar/gku1205

[15] The UniProt Consortium, UniProt: a worldwide hub of protein knowledge, Nucleic Acids Research, Volume 47, Issue D1, 08 January 2019, Pages D506–D515, https://doi.org/10.1093/nar/gky1049

[16] Allan Peter Davis, Cynthia J Grondin, Robin J Johnson, Daniela Sciaky, Jolene Wiegers, Thomas C Wiegers, Carolyn J Mattingly, Comparative Toxicogenomics Database (CTD): update 2021, Nucleic Acids Research, Volume 49, Issue D1, 8 January 2021, Pages D1138–D1143, https://doi.org/10.1093/nar/gkaa891

[17] Orphanet: an online database of rare diseases and orphan drugs. Copyright, INSERM 1997. Available at http://www.orpha.net Accessed (date of access).

[18] ClinGen The Clinical Genome Resource. Heidi L. Rehm, Ph.D., Jonathan S. Berg, M.D., Ph.D., Lisa D. Brooks, Ph.D., Carlos D. Bustamante, Ph.D., James P. Evans, M.D., Ph.D., Melissa J. Landrum, Ph.D., David H. Ledbetter, Ph.D., Donna R. Maglott, Ph.D., Christa Lese Martin, Ph.D., Robert L. Nussbaum, M.D., Sharon E. Plon, M.D., Ph.D., Erin M. Ramos, Ph.D., Stephen T. Sherry, Ph.D., and Michael S. Watson, Ph.D., for ClinGen. N Engl J Med 2015; 372:2235–2242 June 4, 2015 DOI: 10.1056/NEJMsr1406261.

[19] Martin, A. R., Williams, E., Foulger, R. E., Leigh, S., Daugherty, L. C., Niblock, O., Leong, I., Smith, K. R., Gerasimenko, O., Haraldsdottir, E., Thomas, E., Scott, R. H., Baple, E., Tucci, A., Brittain, H., de Burca, A., Ibañez, K., Kasperaviciute, D., Smedley, D., Caulfield, M., … McDonagh, E. M. (2019). PanelApp crowdsources expert knowledge to establish consensus diagnostic gene panels. Nature genetics, 51(11), 1560–1565. https://doi.org/10.1038/s41588-019-0528-2

[20] Tamborero, D., Rubio-Perez, C., Deu-Pons, J. et al. Cancer Genome Interpreter annotates the biological and clinical relevance of tumor alterations. Genome Med 10, 25 (2018). https://doi.org/10.1186/s13073-018-0531-8

[21] Alba Gutiérrez-Sacristán; Solène Grosdidier; Olga Valverde; Marta Torrens; Àlex Bravo; Janet Pinero; Ferran Sanz; Laura I. Furlong “PsyGeNET: a knowledge platform on psychiatric disorders and their genes” Bioinformatics 2015; doi: 10.1093/bioinformatics/btv301

[22] Tian, R., Abarientos, A., Hong, J. et al. Genome-wide CRISPRi/a screens in human neurons link lysosomal failure to ferroptosis. Nat Neurosci (2021). https://doi.org/10.1038/s41593-021-00862-0

[23] Zemojtel, T., Köhler, S., Mackenroth, L., Jäger, M., Hecht, J., Krawitz, P., … & Øien, N. C. (2014). Effective diagnosis of genetic disease by computational phenotype analysis of the disease-associated genome. Science translational medicine, 6(252), 252ra123–252ra123

[24] Kalaria R. Similarities between Alzheimer’s disease and vascular dementia. J Neurol Sci 2002;203–204:29–34.

[25] Bulik-Sullivan B, Finucane HK, Anttila V, et al. An atlas of genetic correlations across human diseases and traits. Nat Genet 2015;47(11):1236–41

[26] Genetic diagnosis in Lafora disease Genotype–phenotype correlations and diagnostic pitfalls H. Lohi, J. Turnbull, X. C. Zhao, S. Pullenayegum, L. Ianzano, M. Yahyaoui, M. A. Mikati, N. P. Quinn, S. Franceschetti, F. Zara, B. A. Minassian Neurology Mar 2007, 68 (13) 996–1001; DOI: 10.1212/01.wnl.0000258561.02248.2f

[27] Köhler S, Schulz MH, Krawitz P, Bauer S, Dölken S, Ott CE, Mundlos C, Horn D, Mundlos S, Robinson PN. Clinical diagnostics in human genetics with semantic similarity searches in ontologies. Am J Hum Genet. 2009; 85(4):457–64

[28] Deng Y, Gao L, Wang B, Guo X. Hposim: an r package for phenotypic similarity measure and enrichment analysis based on the human phenotype ontology. PloS ONE. 2015; 10(2):0115692.

[29] Peng, J., Xue, H., Hui, W., Lu, J., Chen, B., Jiang, Q., … & Wang, Y. (2018). An online tool for measuring and visualizing phenotype similarities using HPO. BMC genomics, 19(6), 89–97.

[30] Weng MP and Liao BY (2017) modPhEA: model organism Phenotype Enrichment Analysis on eukaryotic gene sets. Bioinformatics 33(21):3505–3507

[31] Wang, J., Huang, Q., Liu, Z. P., Wang, Y., Wu, L. Y., Chen, L., & Zhang, X. S. (2011). NOA: a novel Network Ontology Analysis method. Nucleic acids research, 39(13), e87–e87.

[32] Page L, Motwani R, Brin S, Winograd T. The pagerank citation ranking: bringing order to the web. Stanford Digital Libraries Working Paper, 1999. 2009; 9(1):1–14.

[33] Janet Piñero, Juan Manuel Ramírez-Anguita, Josep Saüch-Pitarch, Francesco Ronzano, Emilio Centeno, Ferran Sanz, Laura I Furlong, The DisGeNET knowledge platform for disease genomics: 2019 update, Nucleic Acids Research, Volume 48, Issue D1, 08 January 2020, Pages D845–D855, https://doi.org/10.1093/nar/gkz1021

[34] Braschi B, Denny P, Gray K, Jones T, Seal R, Tweedie S, Yates B, Bruford E. Genenames.org: the HGNC and VGNC resources in 2019. Nucleic Acids Res. 2019 Jan 8;47(D1):D786–D792. PMID:30304474

[35] Winston Chang, Joe Cheng, JJ Allaire, Yihui Xie and Jonathan McPherson (2020). shiny: Web Application Framework for R. R package version 1.5.0. https://CRAN.R-project.org/package=shiny

[36] Niemann, N., & Jankovic, J. (2019). Juvenile parkinsonism: Differential diagnosis, genetics, and treatment. Parkinsonism & related disorders, 67, 74–89.

[37] Breakefield, X. O., Blood, A. J., Li, Y., Hallett, M., Hanson, P. I., & Standaert, D. G. (2008). The pathophysiological basis of dystonias. Nature reviews. Neuroscience, 9(3), 222–234. https://doi.org/10.1038/nrn2337

[38] A. Berardelli, J. C. Rothwell, P. D. Thompson, M. Hallett, Pathophysiology of bradykinesia in Parkinson’s disease, Brain, Volume 124, Issue 11, November 2001, Pages 2131–2146, https://doi.org/10.1093/brain/124.11.2131

[39] Chen, P. H., Wang, R. L., Liou, D. J., & Shaw, J. S. (2013). Gait disorders in Parkinson’s disease: assessment and management. International Journal of Gerontology, 7(4), 189–193.

[40] Hunot, S., & Hirsch, E. C. (2003). Neuroinflammatory processes in Parkinson’s disease. Annals of Neurology: Official Journal of the American Neurological Association and the Child Neurology Society, 53(S3), S49–S60.

[41] O’Keeffe, G. W., & Sullivan, A. M. (2018). Evidence for dopaminergic axonal degeneration as an early pathological process in Parkinson’s disease. Parkinsonism & Related Disorders, 56, 9–15.

[42] Albanese, A., Di Giovanni, M., & Lalli, S. (2019). Dystonia: diagnosis and management. European journal of neurology, 26(1), 5–17.

[43] Brashear, A., Farlow, M. R., Butler, I. J., Kasarskis, E. J., & Dobyns, W. B. (1996). Variable phenotype of rapid-onset dystonia-parkinsonism. Movement disorders: official journal of the Movement Disorder Society, 11(2), 151–156.

[44] Romano, R., Bertolino, A., Gigante, A., Martino, D., Livrea, P., & Defazio, G. (2014). Impaired cognitive functions in adult-onset primary cranial cervical dystonia. Parkinsonism & Related Disorders, 20(2), 162–165.

[45] Furuya, S., Tominaga, K., Miyazaki, F., & Altenmüller, E. (2015). Losing dexterity: patterns of impaired coordination of finger movements in musician’s dystonia. Scientific reports, 5(1), 1–14.

[46] Castagna, A., Frittoli, S., Ferrarin, M., Del Sorbo, F., Romito, L. M., Elia, A. E., & Albanese, A. (2016). Quantitative gait analysis in parkin disease: possible role of dystonia. Movement Disorders, 31(11), 1720–1728.

[47] Booth, H., Hirst, W. D., & Wade-Martins, R. (2017). The Role of Astrocyte Dysfunction in Parkinson’s Disease Pathogenesis. Trends in neurosciences, 40(6), 358–370. https://doi.org/10.1016/j.tins.2017.04.001

[48] Kim, C. Y., Wirth, T., Hubsch, C., Németh, A. H., Okur, V., Anheim, M., … & Chung, W. K. (2020). Early-onset parkinsonism is a manifestation of the PPP2R5D p. E200K mutation. Annals of Neurology, 88(5), 1028–1033

[49] Van Muiswinkel, F. L., De Vos, R. A. I., Bol, J. G. J. M., Andringa, G., Steur, E. J., Ross, D., … & Drukarch, B. (2004). Expression of NAD (P) H: quinone oxidoreductase in the normal and Parkinsonian substantia nigra. Neurobiology of aging, 25(9), 1253–1262.

[50] Zarow, C., Lyness, S. A., Mortimer, J. A., & Chui, H. C. (2003). Neuronal loss is greater in the locus coeruleus than nucleus basalis and substantia nigra in Alzheimer and Parkinson diseases. Archives of neurology, 60(3), 337–341.

[51] Ziemssen, T., & Reichmann, H. (2010). Cardiovascular autonomic dysfunction in Parkinson’s disease. Journal of the neurological sciences, 289(1-2), 74–80.

[52] Aarsland, D., & Kurz, M. W. (2010). The epidemiology of dementia associated with Parkinson disease. Journal of the neurological sciences, 289(1-2), 18–22.

[53] Rosenkranz, K., Williamon, A., Butler, K., Cordivari, C., Lees, A. J., & Rothwell, J. C. (2005). Pathophysiological differences between musician’s dystonia and writer’s cramp. Brain, 128(4), 918–931.

[54] Ibrahim, M. H., Fadhil, A., Ali, S. S., Kader, S. F. A., Khalid, M., Kumar, K., … & Sirsat, J. (2015). Could Dystonia Be Initial Presentation of Corpus Callosum Infarction in Young Age Patients? A Case Report Study. Neuroscience & Medicine, 6(02), 62.

[55] Colosimo C, Pantano P, Calistri V,et alDiffusion tensor imaging in primary cervical dystoniaJournal of Neurology, Neurosurgery & Psychiatry 2005;76:1591–1593.

[56] Schneider, S. A., Lang, A. E., Moro, E., Bader, B., Danek, A., & Bhatia, K. P. (2010). Characteristic head drops and axial extension in advanced chorea-acanthocytosis. Movement disorders, 25(10), 1487–1491.

[57] Gorman, K. M., Meyer, E., & Kurian, M. A. (2018). Review of the phenotype of early-onset generalised progressive dystonia due to mutations in KMT2B. European Journal of Paediatric Neurology, 22(2), 245–256.

[58] Lohmann, K., & Klein, C. (2017). Update on the genetics of dystonia. Current neurology and neuroscience reports, 17(3), 26.

[59] Groffen, A. J., Klapwijk, T., van Rootselaar, A. F., Groen, J. L., & Tijssen, M. A. (2013). Genetic and phenotypic heterogeneity in sporadic and familial forms of paroxysmal dyskinesia. Journal of neurology, 260(1), 93–99. https://doi.org/10.1007/s00415-012-6592-5

[60] Botía, J. A., Guelfi, S., Zhang, D., D’Sa, K., Reynolds, R., Onah, D., … & Ryten, M. (2018). G2P: Using machine learning to understand and predict genes causing rare neurological disorders. bioRxiv, 288845.

[61] Stafstrom, C. E., & Carmant, L. (2015). Seizures and epilepsy: an overview for neuroscientists. Cold Spring Harbor perspectives in medicine, 5(6), a022426.

[62] Ishiura, H., Doi, K., Mitsui, J., Yoshimura, J., Matsukawa, M. K., Fujiyama, A., … & Tsuji, S. (2018). Expansions of intronic TTTCA and TTTTA repeats in benign adult familial myoclonic epilepsy. Nature genetics, 50(4), 581–590.

[63] Trinka, E., Höfler, J., & Zerbs, A. (2012). Causes of status epilepticus. Epilepsia, 53, 127–138.

[64] Abdel-Salam, G. M., Halász, A. A., & Czeizel, A. E. (2000). Association of epilepsy with different groups of microcephaly. Developmental Medicine & Child Neurology, 42(11), 760–767.

[65] Carvalho, M. D. C., Ximenes, R. A., Montarroyos, U. R., da Silva, P. F., Andrade-Valença, L. P., Eickmann, S. H., … & Microcephaly Epidemic Research Group. (2020). Early epilepsy in children with Zika-related microcephaly in a cohort in Recife, Brazil: Characteristics, electroencephalographic findings, and treatment response. Epilepsia, 61(3), 509–518.

[66] Ricobaraza, A., Mora-Jimenez, L., Puerta, E. et al. Epilepsy and neuropsychiatric comorbidities in mice carrying a recurrent Dravet syndrome SCN1A missense mutation. Sci Rep 9, 14172 (2019). https://doi.org/10.1038/s41598-019-50627-w

[67] Parisi, P., Moavero, R., Verrotti, A., & Curatolo, P. (2010). Attention deficit hyperactivity disorder in children with epilepsy. Brain and Development, 32(1), 10–16.

[68] Lee, B. H., Smith, T., & Paciorkowski, A. R. (2015). Autism spectrum disorder and epilepsy: disorders with a shared biology. Epilepsy & Behavior, 47, 191–201.

[69] Tuchman, R., & Rapin, I. (2002). Epilepsy in autism. The Lancet Neurology, 1(6), 352–358.

[70] Epi25 Collaborative. Ultra-Rare Genetic Variation in the Epilepsies: A Whole-Exome Sequencing Study of 17,606 Individuals. Am J Hum Genet. 2019;105(2):267–282. https://doi.org/10.1016/j.ajhg.2019.05.020

